# Mitotic H3K9ac is controlled by phase-specific activity of HDAC2, HDAC3 and SIRT1

**DOI:** 10.1101/2021.03.08.434337

**Authors:** Shashi Gandhi, Raizy Mitterhoff, Rachel Rapoport, Marganit Farago, Avraham Greenberg, Lauren Hodge, Sharon Eden, Christopher Benner, Alon Goran, Itamar Simon

**Affiliations:** Department of Microbiology and Molecular Genetics, Institute of Medical Research Israel-Canada, Faculty of Medicine, The Hebrew University, Jerusalem 91120, Israel; Department of Medicine, University of California, San Diego, La Jolla, California 92093, USA

**Keywords:** HDACs, histone modifications, Chromatin, Mitosis

## Abstract

Histone acetylation levels are reduced during mitosis. To study the mitotic regulation of H3K9ac we used an array of inhibitors targeting specific histone deacetylases. We evaluated the involvement of the targeted enzymes in regulating H3K9ac during all mitotic stages by immunofluorescence and immunoblots. We identified HDAC2, HDAC3 and SIRT1 as modulators of H3K9ac mitotic levels. HDAC2 inhibition increased H3K9ac levels in prophase, whereas HDAC3 or SIRT1 inhibition increased H3K9ac levels in metaphase. Next, we performed ChIP-seq on mitotic-arrested cells following targeted inhibition of these histone deacetylases. We found that both HDAC2 and HDAC3 have a similar impact on H3K9ac, and inhibiting either of these two HDACs substantially increases the levels of this histone acetylation in promoters, enhancers and insulators. Altogether, our results support a model in which H3K9 deacetylation is a stepwise process – at prophase HDAC2 modulates most transcription-associated H3K9ac-marked loci and at metaphase HDAC3 maintains the reduced acetylation, whereas SIRT1 potentially regulates H3K9ac by impacting HAT activity.

**Summary blurb:** Combination of immunofluorescence, western blot and ChIP-seq revealed the interplay between HDAC2, HDAC3 and SIRT1 in H3K9 deacetylation during mitosis of mammalian cells.

## Introduction

The eukaryotic cell cycle is composed of many cellular events. Throughout interphase the genome and other cellular components are duplicated, and during mitosis the cell is divided into two daughter cells. The relatively short phase of mitosis is tightly regulated as it is critical to ensure that the cellular identity and function are correctly relayed to daughter cells (McIntosh & Hays 2016, Palozola et al 2019). The major molecular changes occurring during mitosis include a striking decrease in transcription (Prescott & Bender 1962), condensation of the chromatin (Antonin & Neumann 2016), nuclear envelope breakdown (Robbins & Gonatas 1964), and loss of long-range intra-chromosomal interactions (Naumova et al. 2013; Dileep et al. 2015; Stevens et al. 2017).

We have conducted a comprehensive study to assess the changes in histone modifications during mitosis (Javasky et al 2018). Our observations, together with other studies (Behera et al 2019, Ginno et al 2018, Hsiung et al 2015, Kang et al 2020, Kelly et al 2010, Kruhlak et al 2001, Liang et al 2015, Liu et al 2017, McManus & Hendzel 2006, Pelham-Webb et al 2021, Van Hooser et al 1998) support the following notions: (***i***) the global epigenetic landscape is preserved during mitosis; (***ii***) the mitosis phase encompasses global reduction in histone acetylation; and (***iii***) there is a dramatic increase in histone phosphorylation.

What causes histone deacetylation during mitosis? Histone acetylation is carried out by HATs (Lee & Workman 2007) and is removed by HDACs (Haberland et al 2009). HDACs encompass a diverse set of deacetylases that are involved in the regulation of the acetylation levels of histones as well as many other proteins. There are 18 HDACs in mammals that are classified into four major classes based on their homology to yeast HDACs. This categorization includes HDAC1-3 and HDAC8 in Class I, HDAC4-10 in Class II, the sirtuins SIRT1-7 in Class III and HDAC11 in Class IV (Seto & Yoshida 2014). While some evidence has linked specific HDACs to mitosis, the exact HDAC that conducts the mitotic deacetylations is not well characterized. HDAC3 was suggested to play a key role in human mitosis, since knocking it down in human HeLa cells affects mitosis histone deacetylation (Li et al 2006). In yeast, the Hst2p histone deacetylase (SIR2 homologue), was shown to be responsible for the deacetylation of H4K16 during mitosis (Wilkins et al 2014). In addition, the mitosis-specific deacetylation of histones could be carried out by a change in the balance between HATs and HDACs. Evidence supporting this notion was provided by a mass spectrometry-based study that determined the changes in DNA-associated proteins between various cell cycle stages and demonstrated a general retention of HDACs and depletion of HATs in mitosis (Ginno et al 2018).

Here, we employed an array of small molecule inhibitors, each one targeting a different set of HDACs or a specific HDAC (**Table 1**). We then measured the impact of these perturbations on deacetylation of H3K9 by immunofluorescence, immunoblots and ChIP-seq. We observed that three histone deacetylases, namely – HDAC2, HDAC3 and SIRT1 – are involved in H3K9 deacetylation. Further dissection of the roles each histone deacetylase plays revealed that HDAC2 initially deacetylates H3K9 at prophase whereas HDAC3 activity is only detected later during metaphase. While we noticed that SIRT1 is absent from the mitotic chromatin, we observed that this histone deacetylase potentially affects H3K9 acetylation indirectly through the modulation of mitotic HAT activity. Taken together, our results provide insight into the biochemical pathways involved in histone deacetylation during mitosis and pave the way for future studies aimed at deciphering the role the deacetylation process plays in regulation of mitotic gene expression and open chromatin.

**Table 1.** HDACs inhibitors and their specificty.

| Name | Specificity | Concentration used | References |
| --- | --- | --- | --- |
| TSA | Pan-HDACs | 150 nM | (Khan et al 2008, Lobera et al 2013) |
| CHIDAMIDE | HDAC1,2,3,10 | 10 μM | (Ning et al 2012) |
| RGFP966 | HDAC3 | 40 μM | (Malvaez et al 2013) |
| CAY10683 | HDAC2 and HDAC6 | 1 μM | (Jaju Bhattad et al 2020, Pavlik et al 2013) |
| MS-275 | HDAC1,2,3 | 1 μM | (Khan et al 2008) |
| NICOTINAMIDE | Pan-sirtuins | 10 mM | (Hu et al 2014) |
| EX-527 | SIRT1 | 5nM | (Hu et al 2014) |

## Results

### Identification of the deacetylases involved in modulating H3K9ac during mitosis

To study the changes in histone acetylation during mitosis, we focused on H3K9ac, which is reduced approximately by 2.7-fold during mitosis in HeLa-S3 cells (Javasky et al 2018). To better identify the mitotic stage in which deacetylation takes place, we enriched HeLa-S3 cells for mitotic cells by releasing the cells from a double thymidine block for 8.5 hours. This resulted in primarily G2/M cells with approximately 35% of the cells in mitosis, most of them in the metaphase stage (**Fig. 1a** and **Supplementary Figure S1**). Immunofluorescence of H3K9ac at different mitotic stages revealed that the H3K9ac mark declines at prophase, reaches a minimal level at metaphase, remains low at anaphase and gradually increases at later stages of mitosis (**Fig. 1b**).

**Figure 1.**
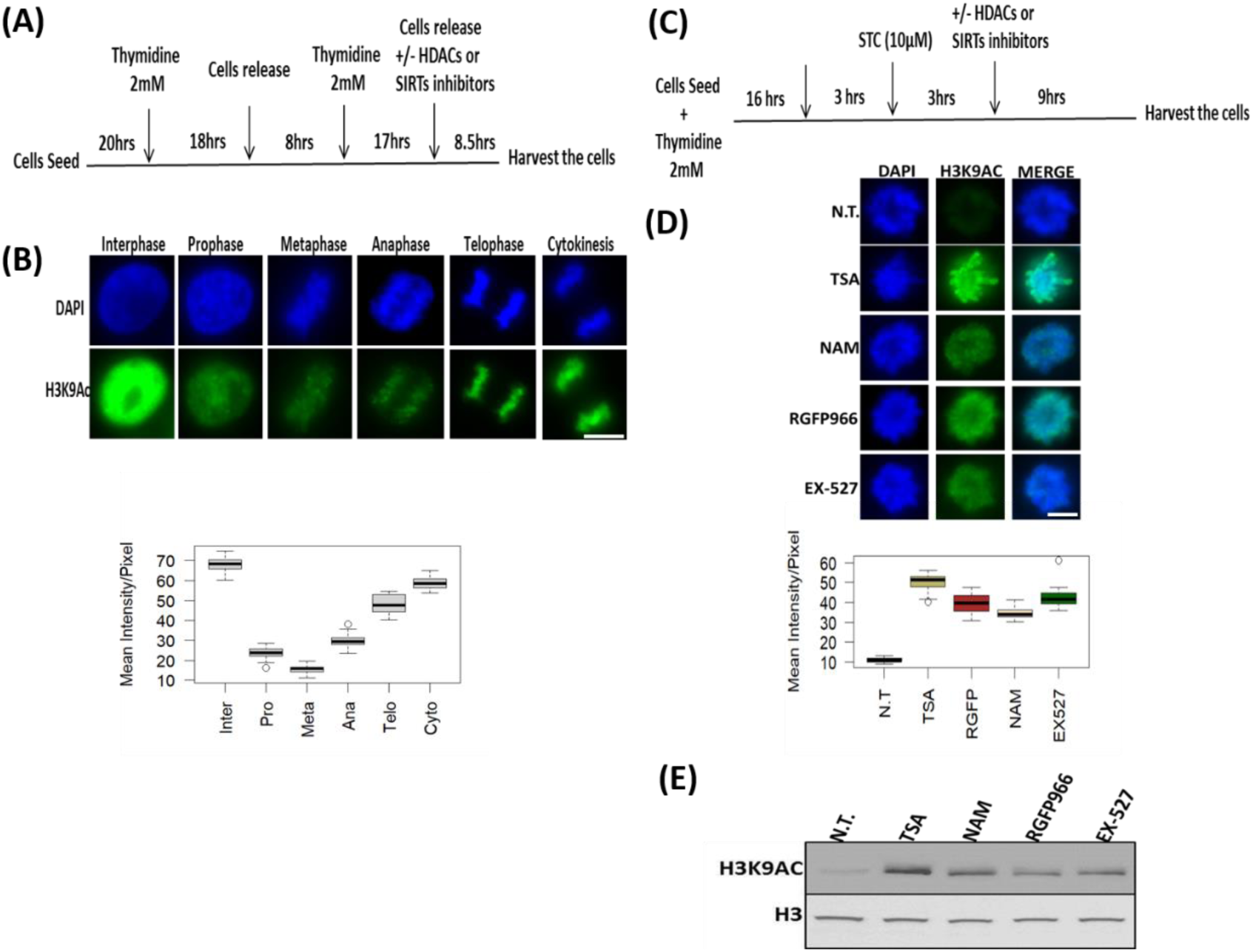
H3K9ac levels during mitosis. **(a)** Schematic representation of the synchronization strategy by double thymidine block method. **(b)** Immunofluorescence of HeLa-S3 cells enriched for mitotic stages stained for H3K9ac (green), and DNA (blue). Representative pictures from the major mitosis stages are shown. Quantification of the immunofluorescence results presented below the pictures. Intensity values represent the mean ± SEM of at least 20 cells for every stage. **(c)** Schematic representation of the synchronization strategy by kinesin-5 inhibitor S-trityl-L-cysteine (STC). **(d)** Immunofluorescence of STC arrested HeLa-S3 cells stained for DNA (blue) and H3K9ac (green). The cells were either not treated (N.T.) or treated with TSA (150nM); Nicotinamide (NAM, 10mM); RGFP966 (40μM); EX-527 (5nM). Quantification of the immunofluorescence results are shown below the pictures. Intensity values represent the mean ± SEM of at least 25 cells for every condition. All treatments significantly (P<10^−23^; one sided ttest) increased H3K9ac levels **(e)** Immunoblot showing the H3K9ac signal for synchronized HeLa-S3 cells treated with the indicated inhibitors (same concentration as in **d**), total histone H3 serves as a loading control. Scale bars:5μm

To identify the HDACs involved in the decrease of H3K9ac levels in metaphase, we arrested the cells at a metaphase-like stage with monoastral spindles using the kinesin 5 inhibitor STC (Skoufias et al 2006). We added various HDAC inhibitors for 9 hours (**Fig. 1c**; **Table 1**) and measured H3K9ac levels by immunofluorescence (**Fig. 1d**). We found that treatment with either the pan-HDAC inhibitor trichostatin A (TSA) (Khan et al 2008, Lobera et al 2013) or the pan-sirtuin inhibitor nicotinamide (NAM) (Hu et al 2014) induces a significant increase (P<10^−23^; one sided t test) in H3K9ac levels.

We further studied the deacetylation process by using specific inhibitors for particular histone deacetylases. Building on the results of previous studies linking HDAC3 and SIRT1 to mitosis (Fatoba & Okorokov 2011, Li et al 2006), we decided to first investigate the involvement of these histone deacetylases. We observed that the HDAC3-specific inhibitor RGFP966 (Malvaez et al 2013) and the SIRT1-specific inhibitor EX-527 (Hu et al 2014) induce an increase in mitotic H3K9ac that is similar to the acetylation levels following inhibition with pan-HDAC or pan-sirtuin inhibitors (**Fig. 1d**). Similar results were obtained by immunoblots (**Fig. 1e**).

The deacetylation of H3K9 initiated at prophase (**Fig. 1b**); consequently, it cannot be fully studied by STC synchronization that arrests cells at a metaphase-like stage. We therefore targeted double-thymidine synchronized cells with the small-molecule inhibitors. The various inhibitors (**Table 1**) were used upon release from the second thymidine block, and the level of H3K9ac was assessed 8.5 hours later, a time-point that is maximally enriched for cells from all mitotic stages and cytokinesis (**Fig. 1a** and **Supplementary Figure S1**). Cells were binned according to the different mitosis stages (prophase, metaphase, anaphase, and telophase) and cytokinesis, and H3K9ac immunofluorescence intensity was measured separately for each stage (**Fig. 2a**). We found that the general HDAC inhibitor TSA (Khan et al 2008, Lobera et al 2013) affects H3K9ac at prophase, while the pan-sirtuin inhibitor NAM (Hu et al 2014) induces an increase in acetylation levels only in metaphase.

**Figure 2.**
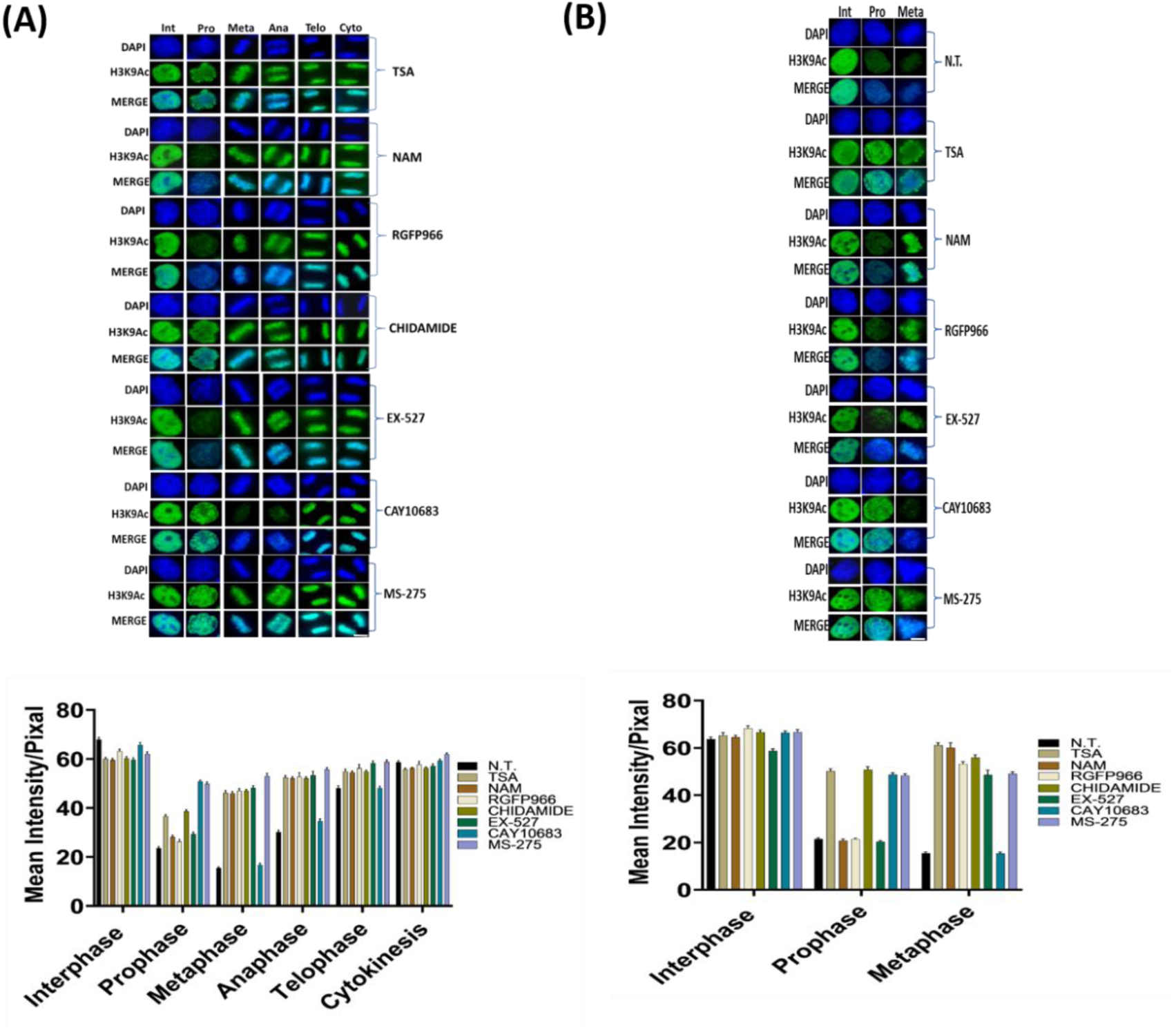
HDAC inhibition shows involvement of specific HDACs in modulating the dynamics of H3K9ac during mitosis. (a)Immunofluorescence of HeLa-S3 cells enriched for mitotic stages stained for H3K9ac (green), and DNA (blue). Representative pictures from the major mitosis stages are shown. The cells were either not treated (N.T.) or treated with TSA (150nM); Nicotinamide (NAM, 10mM); RGFP966 (40μM); Chidamide (10μM); EX-527 (5nM); CAY10683 (1μM); MS-275 (1μM). Below, quantification of the immunofluorescence results. Intensity values represent the mean ± SEM of at least 20 cells for every stage. **(b)** Immunofluorescence of MEFs cells enriched for mitotic stages stained for H3K9ac (green), and DNA (blue). Representative pictures from the Interphase, Prophase and Metaphase stages are shown. The cells were either not treated (N.T.) or treated with TSA (150nM); Nicotinamide (NAM, 10mM); RGFP966 (40μM); Chidamide (10μM); EX-527 (5nM); CAY10683 (1μM); MS-275 (1μM). Below, quantification of the immunofluorescence results. Intensity values represent the mean ± SEM of at least 20 cells for every stage. All treatments (both in HeLa-S3 and MEF cells) beside CAY10863 significantly affect H3K9ac levels at metaphase (P<10^−15^, FDR corrected t-test). Scale bars:5μm. for a boxplot version of the graphs, see **supplementary figure S3**.

To identify the specific enzyme or enzymes that are involved in mitotic deacetylation, we used small molecules that selectively target a single or a concrete subset of histone deacetylases (**Table 1**). In particular, we observed that treatment with RGFP966, an HDAC3-specific inhibitor (Malvaez et al 2013), affects H3K9ac levels only in metaphase, while treatment with Chidamide that selectively inhibits HDAC1, HDAC2, HDAC3 and HDAC10 (Ning et al 2012), shows very similar results to TSA. These results suggest that the deacetylation we observed at prophase is probably carried out by either HDAC1, HDAC2 or HDAC10, given the differential impact of inhibiting HDAC3 alone by RGFP966.

To determine which of the three HDACs (HDAC1, HDAC2 or HDAC10) is modulating H3K9ac during prophase, we used two specific inhibitors – MS-275 that specifically targets HDAC1, HDAC2 and HDAC3 (Khan et al 2008, Tatamiya et al 2004) and CAY10683, an HDAC2 inhibitor (Jaju Bhattad et al 2020, Pavlik et al 2013). Following the specific inhibition, we measured H3K9ac levels in cells captured at different stages of mitosis (**Fig. 2a**). We observed that both MS-275 and CAY10683 induced an increase in H3K9 acetylation levels in prophase. However, only MS-275 induced an increase in H3K9 acetylation levels in metaphase as well. This specific pattern suggests that HDAC2 (targeted by CAY10683 and MS-275) deacetylates H3K9ac during prophase, while HDAC3 (targeted by RGFP966 and MS-275), which is not active in prophase, maintains the low acetylation levels from metaphase through telophase. Repeating the measurements with a larger sample of cells revealed that the CAY10683 inhibitor was able to induce a small increase in H3K9ac in metaphase as well (**Supplementary Figure S2**).

As mentioned above, the pan-sirtuin inhibitor, nicotinamide (NAM) induces a significant increase in H3K9ac levels as well. To study the involvement of the NAM-target sirtuins in H3K9 deacetylation in mitosis, we used the SIRT1 specific inhibitor, EX-527 (Hu et al 2014). The effect of SIRT1 inhibition was very similar to that of pan-sirtuin inhibition, suggesting that among the sirtuins, SIRT1 is the main mitotic H3K9 deacetylase.

We then evaluated whether the results we obtained in HeLa-S3 can be reproduced in another system by repeating the main experiments in mouse embryonic fibroblasts (MEFs). To this end, we synchronized the MEFs with a double thymidine block and measured H3K9ac levels by immunofluorescence 8.5 hours after release from the second block. We focused on prophase and metaphase in the MEFs and observed high similarity to the results obtained in the human cell line (HeLa-S3). Specifically, treatment of MEFs with either TSA, Chidamide or MS-257 induced an increase in H3K9ac levels in both prophase and metaphase (**Fig. 2b**). On the other hand, the increased H3K9ac level in MEFs was seen only in metaphase when treated with NAM, EX-527, or RGFP966, and only in prophase when treated with CAY10683 (**Fig. 2b**).

Taken together, these results suggest that mitosis-associated H3K9 deacetylation is a two-stage process – HDAC2 performs deacetylation during prophase, and HDAC3 and SIRT1 subsequently maintain low acetylation levels in metaphase.

### Variations in the mitotic chromatin association of key histone deacetylases and histone acetyltransferases

Our finding that the deacetylation process is dependent on HDAC2, HDAC3 and SIRT1, suggests that these deacetylases are associated with the mitotic chromosomes. We carried out immunofluorescence experiments targeting key HDACs and sirtuins to evaluate their mitotic localization. Using our double thymidine synchronization scheme (**Fig. 1a**), we observed that among the HDACs tested, mainly HDAC3 was retained on the mitotic chromosomes during all mitotic stages. On the other hand, HDAC1 and HDAC2 appeared to be associated with the mitotic chromosomes at lower levels during metaphase (**Fig. 3a**). Surprisingly, even though we observed that inhibition of SIRT1 brings about an increase in H3K9ac during metaphase, neither SIRT1 nor the other sirtuins we tested (SIRT2, SIRT6 and SIRT7) showed strong binding to the chromatin in metaphase. While we observed low binding of SIRT1 during prophase, there was a reduction in its chromatin localization starting at metaphase for the duration of mitosis (**Fig. 3b**).

**Figure 3.**
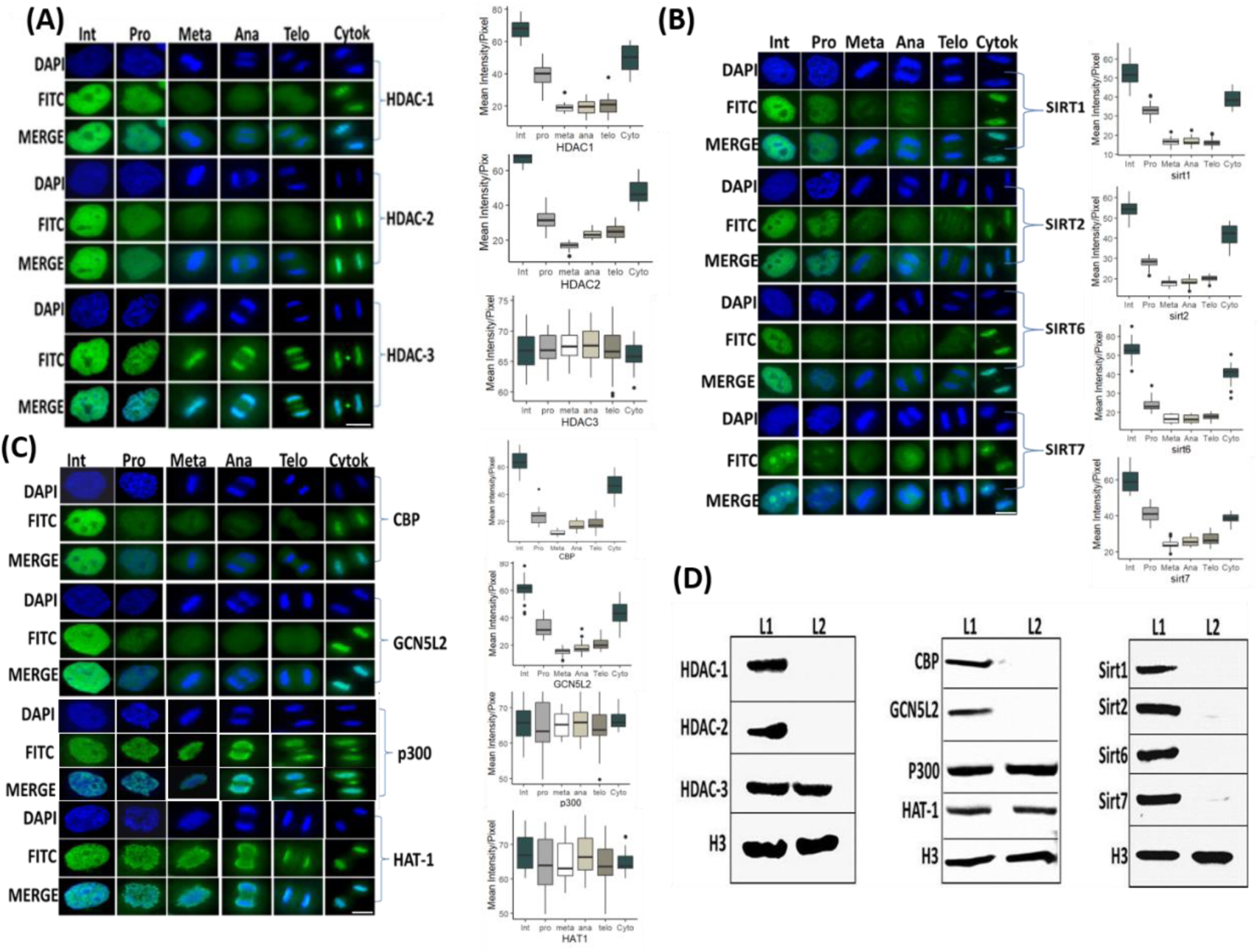
Identification of the chromatin localization patterns of key HATs and HDACs during mitosis. Immunofluorescence of HeLa-S3 cells enriched for mitotic stages, stained for DNA (blue) and for various HDACs **(a)**, sirtuins **(b)** and HATs **(c)** (green). Representative pictures from the major mitosis stages and interphase are shown, along with a quantification of at least 20 cells from each stage. Intensity values represent the mean ± SEM. Note that only HDAC3, P300 and HAT-1 are retained on the mitotic chromatin along all mitotic stages, whereas the other proteins show partial retention at prophase. All factors (beside HDAC3, P300 and HAT1) show significant reduction at metaphase (P<10^−16^, FDR corrected t-test) **(d)** Immunoblots showing the abundance of the indicated chromatin modifiers in the chromatin fraction of interphase (L1) and mitotic (L2) HeLa-S3 cells. Total histone H3 serves as loading control. Scale bars:5μm.

We performed similar experiments with several histone acetyl transferases (HATs) and, observed that HAT1 and EP300, but not CBP and GCNL2, retained their binding to the mitotic chromosomes. Interestingly, both HAT1 and EP300 seem to be more spread out on the metaphase chromsomes than HDAC3 (**Fig. 3c**). Next, we validated these results by immunoblotting of chromatin from metaphase (STC arrested) and interphase cells. In a similar manner to the immunofluorescence experiments, we detected a signal only for HDAC3, HAT1 and EP300 from the metaphase chromatin fraction. On the other hand, using mitotic chromatin we could not identify a signal for HDAC1, HDAC2, SIRT1, 2, 6 and 7 or two of the HATs – CBP and GCNL2 (**Fig. 3d**). Together, the localization of HDAC2 to mitotic chromatin at prophase and the binding of HDAC3 throughout mitosis are in line with the two-stage model presented above, while the binding of SIRT1 at prophase was unexpected.

### SIRT1 potentially impacts H3K9ac levels indirectly by modulating HAT activity

We considered two hypotheses to reconcile our seemingly contradictory observations, namely, that SIRT1 inhibition impacts H3K9ac levels during mitosis whilst also showing low binding to chromatin during mitosis. One option is that SIRT1 directly deacetylates H3K9 prior to metaphase, before it detaches from the mitotic chromosomes. However, we ruled out this possibility since inhibition of SIRT1 by EX-527 does not appear to impact the prophase levels of H3K9ac (**Fig. 2a**). The second option we considered is that SIRT1 indirectly impacts the H3K9ac levels at metaphase by modulating the activity of other histone deacetylases or acetyl transferases.

We developed an *in vitro* assay to study the impact of SIRT1 on histone acetyl transferase activity in metaphase (**Fig. 4**). We isolated protein extracts from STC-arrested HeLa-S3 cells either treated or untreated with the SIRT1 inhibitor EX-525. The extracts were used to acetylate His-tagged histone H3 peptide, and HAT activity was studied via two approaches. First, we used a colorimetric assay that measures the amount of Co-A released in the acetylation reaction via the DTNB absorbance at 412nm (Foyn et al 2017). Additionally, we immunoprecipitated the His-tagged histone H3 peptide using nickel beads and evaluated the acetylation level of this H3 peptide by immunoblot using an H3K9ac antibody. Both assays revealed that HAT activity was increased in extracts made from cells treated with the SIRT1 inhibitor Ex-527 (**Fig. 4a-b**), suggesting that SIRT1 represses the cellular HAT activity during the metaphase stage.

**Figure 4.**
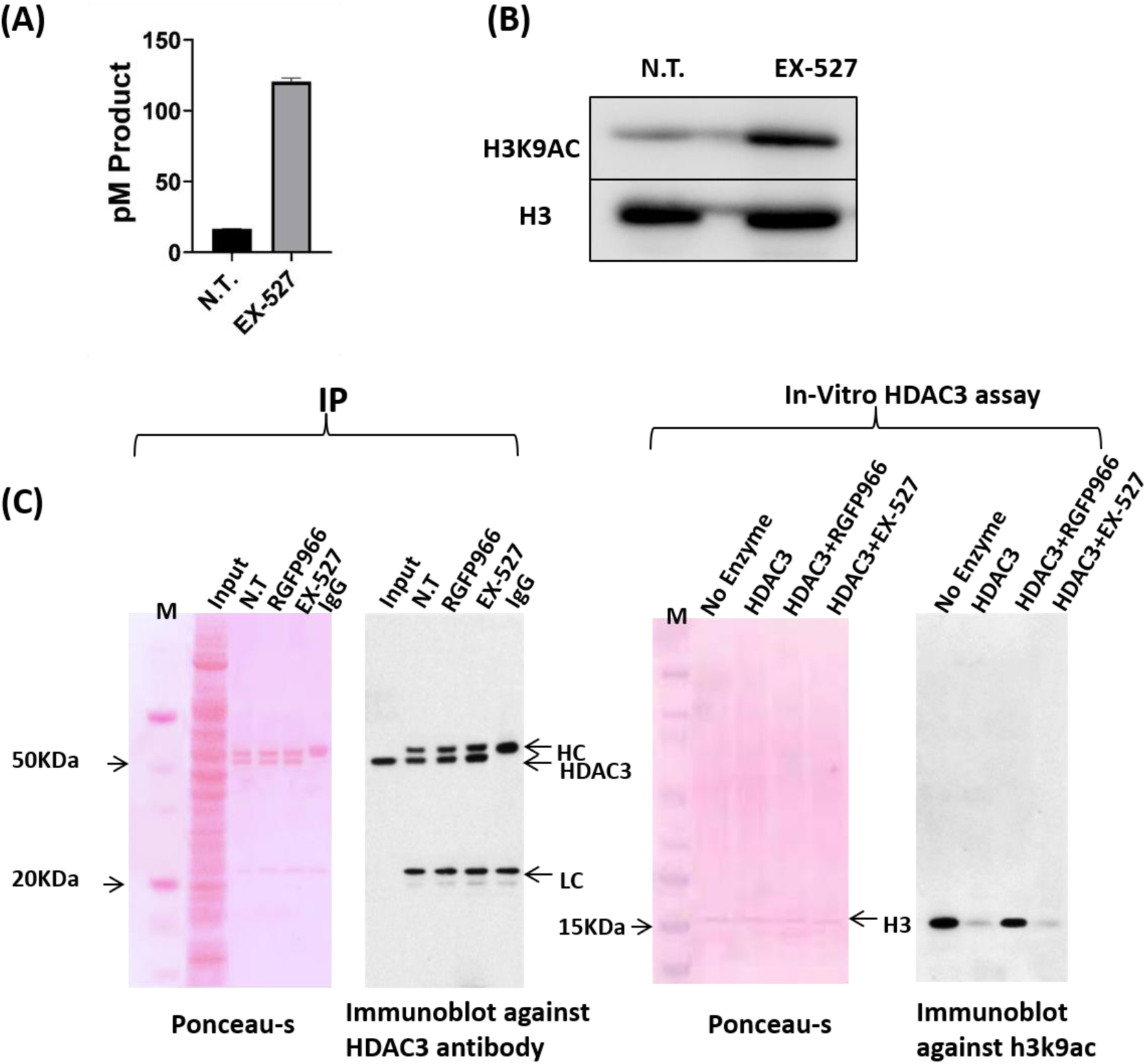
An in vitro assay supports an indirect modulation H3K9ac levels by impacting HAT activity. (**a**) HAT activity colorimetric assay results (mean +/-standard error, N=2) for HeLa-S3 cells either not treated (N.T.) or treated with EX-527 (5nM). **(b)** Immunoblot analysis showing H3K9 acetylation on His-tagged H3 after HAT activity of HeLa mitotic chromatin either not treated (N.T.) or treated with EX-527 (5nM) on His-tagged H3. Total histone H3 serves as loading control. **(c)** Immunoprecipitation of HDAC3 tested by immunoblotting. HDAC3 antibodies and protein A magnetic beads were used to immunoprecipitate (IP) HDAC3 complexes from 1mg of HeLa mitotic extract either not treated (N.T.) or treated with RGFP966 (40μM) or EX-527 (5nM). (i) Showing ponceau S staining of HDAC3 immunocomplex; (ii) showing immunoblot using HDAC3 antibodies developed by ECL method; (iii) showing ponceau S staining of HDAC3 activity assay reaction subjected to immunoblot and (iv) showing immunoblot analysis of immunoprecipitated HDAC3 activity on acetylated his-tagged H3 protein.

Next, to test whether SIRT1 is impacting a histone deacetylase we focused on HDAC3. Our reasoning was that HDAC3 was the only histone deacetylase we detected on the metaphase chromatin (**Fig. 3**), and that HDAC3 is highly involved in regulating metaphase H3K9ac (**Fig. 2a**). To evaluate the impact of SIRT1 on HDAC3 activity in metaphase, we immunoprecipitated HDAC3 from STC-arrested mitotic cells treated or untreated with EX-527. We then used acetylated His-tagged histone H3 peptide and measured H3K9ac levels by immunoblot. We observed that inhibition of SIRT1 does not impact the deacetylase activity of HDAC3 towards H3K9ac (**Fig. 4c**). Taken together, we show that SIRT1 appears to indirectly affect H3K9ac levels by potentially modulating mitotic HAT activity. This result is in line with the observation that SIRT1 is mostly detached from the chromosomes during mitosis (**Fig. 3b**).

### HDAC2 and HDAC3 activity regulate both common and unique genomic loci

The results described above suggest that deacetylation of H3K9 is conducted by HDAC2 in prophase and by HDAC3 in metaphase, with SIRT1 indirectly involved in an additional reduction in H3K9ac during metaphase via modulation of HAT activity.

However, these results were obtained by immunofluorescence and immunoblots, and thus do not provide information about the regulation of histone acetylation at a genomic level by these deacetylases.

To determine genomic context of H3K9 deacetylation, we performed ChIP-seq for H3K9ac on STC-arrested cells, either treated or untreated with CAY10683 (HDAC2 inhibitor), RGFP966 (HDAC3 inhibitor), MS-275 (HDAC2 and HDAC3 inhibitor) and EX-527 (SIRT1 inhibitor) (**Table 1**). To quantitatively compare H3K9ac enrichment between the ChIP-seq experiments we used the ChIP-Rx approach (Orlando et al 2014) which normalizes the efficiency of ChIP by adding the same amount of chromatin from a distinct organism to every reaction. To this end, we added chicken chromatin (harvested from 50,000 untreated and unsynchronized DT-40 cells) to each ChIP-seq reaction. H3K9ac is expected to be enriched in promoters, enhancers and insulators as we observed previously (Javasky et al 2018, Zhou et al 2011). As expected, the chicken chromatin presented a strong enrichment of H3K9ac at these genomic loci, thus validating our ChIP-seq conditions. We used the chicken promoter occupancy to normalize for the differences in ChIP efficiency between the samples (**Methods, Fig. 5a** and **Supplementary Figure S4)**.

**Figure 5.**
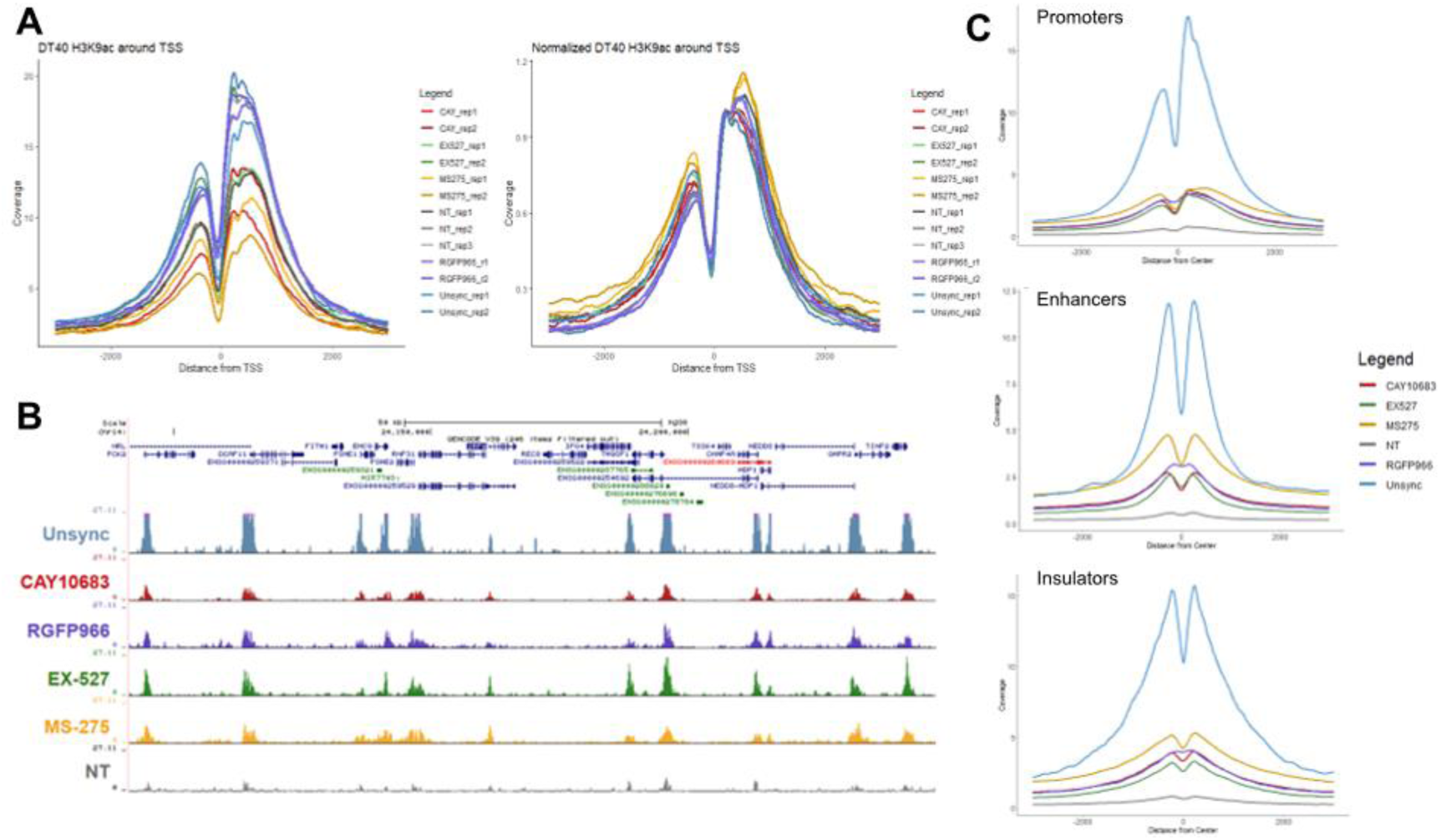
Detection of genomic patterns associated with the activity of HDAC2 and HDAC3. (a)Metagene plots showing H3K9ac promoter occupancy in chicken DT-40 cells before (left) and after (right) normalization. (**b**) Genomic viewer (IGV) tracks representing the H3K9ac ChIP-seq enrichment for the indicated conditions. (**c**) Metagene plots showing H3K9ac occupancy around promoters, enhancers and insulators in HeLa-S3 cells. The data were normalized using the DT-40 promoter data. Similar results were obtained in a biological repeat (**Supplementary Figure S5)**.

We used the normalized signal and compared H3K9ac occupancy at promoters, enhancers and insulators in the different conditions (**Fig. 5b, c**). In line with previous results by us and others (Javasky et al 2018, Palozola et al 2019), H3K9ac levels were high in unsynchronized (interphase) cells, whereas the acetylation declines during mitosis in untreated cells. Interestingly, all of the inhibitors we used had a similar impact on the levels of H3K9ac, with MS-275 (which inhibits both HDAC2 and HDAC3) having a slightly stronger effect when compared to CAY10683 (which inhibits only HDAC2) or RGFP966 (which inhibits only HDAC3). Of note, while the specific inhibition of HDACs in mitotic cells has demonstrated a strong increase in H3K9ac levels, the abundance of the signal was still lower than observed in unsynchronized, interphase cells. This potentially suggests that the regulation of histone deacetylation in mitosis involves the combination of multiple HDACs as well as a reduction in HAT activities.

## Discussion

Multiple studies have provided evidence for the presence of a histone deacetylation process during mitosis (Palozola et al 2019). However, to our knowledge, the mechanism underlying this process has not been previously studied in detail. Here, we combined molecular and genomic techniques and an array of small molecules to identify the involvement of specific histone deacetylases during mitosis. Using this approach, we were able to detect three regulators that are playing a key role in the mitosis-associated deacetylation of H3K9ac – HDAC2, HDAC3 and SIRT1 (**Fig. 2**). In particular, we found that HDAC2 deacetylates H3K9ac in prophase and HDAC3 deacetylates this histone modification during metaphase. While we observed that SIRT1 becomes detached from the chromatin during mitosis, we provide *in vitro* evidence that supports involvement of this sirtuin in impacting the levels of H3K9ac via repression of mitotic HAT activity.

Although inhibition of HDAC2 increases H3K9ac levels at prophase, the acetylation decreases again in metaphase, most probably due to the activity of HDAC3. Indeed, when we inhibited both HDAC2 and HDAC3 we observed an increase in H3K9ac levels throughout mitosis. On the other hand, HDAC3 inhibition did not affect H3K9ac levels at prophase (probably due to the activity of HDAC2) but induced an increase in H3K9ac levels at metaphase. By performing ChIP-seq on STC-arrested cells treated with various HDAC inhibitors we identified the genomic locations of activity for each of these HDACs. We found that inhibition of either HDAC2 or HDAC3 has a similar effect on promoters, enhancers and insulators (**Fig. 5c**). Yet, there are certain genomic regions that appear to be preferably deacetylated by HDAC3 (**Fig. S5**). These results are in line with our immunofluorescence results (**Fig 2a**) showing that HDAC3 has a stronger impact on H3K9ac levels during metaphase (most of the cells during an STC arrest are in metaphase).

Under normal growth conditions, increases in H3K9ac start at anaphase (**Fig. 1b**). We observed that two HATs (HAT1 and EP300) remain associated with the mitotic chromosomes at all mitosis stages (**Fig. 3**), however, HAT activity during metaphase appears to be reduced indirectly by SIRT1 (**Fig. 4a-b**). This suggests that H3K9ac increases are delayed to anaphase due to the modulation of HAT activity by post translation modification (PTM). Indeed, SIRT1 inhibition causes an increase in H3K9ac levels already at metaphase (**Fig. 2**). Modulation of HAT activity by PTMs is a well-documented phenomenon. Many of the HATs are activated by PTMs (Pavlik et al 2013) and are frequently activated by auto-acetylation (McCullough & Marmorstein 2016). Thus, it is reasonable to assume that HAT deacetylation may serve as a tool to partially repress the activity of key HATs during mitosis. Indeed, it was shown that EP300 is repressed by SIRT1-mediated deacetylation at lysine residues 1020/1024 (Bouras et al 2005). Our observation is in line with these observations and suggests a physiological role for the regulation of HATs by SIRT1 at mitosis.

Our results suggest that the low levels of H3K9ac during mitosis are achieved by a combination of two HDACs and by the modulation of HAT activity by a third histone deacetylase. Indeed, it was shown previously that the sole inhibition of class I and II HDACs by sodium butyrate is not sufficient to achieve full acetylation during mitosis due to the decreased ability of HATs to act on histones in mitotic chromatin (Patzlaff et al 2010). It has been shown that RNA polymerase and many of the transcription factors fall off the mitotic chromatin (Kadauke & Blobel 2013, Martinez-Balbas et al 1995, Parsons & Spencer 1997). Yet, the assumption that almost all transcription regulators are displaced from the chromatin during mitosis was recently challenged by a mass-spectrometry based study that showed a large-scale retention of TFs and preinitiation complex members on the mitotic chromatin (Ginno et al 2018). Indeed, our results regarding the mitotic localization of several histone acetylases and deacetylases support the more complex picture of partial retention of transcription regulators on the mitotic chromatin (**Fig. 3**). We found that HDAC1 and HDAC2 fall off the chromatin only at metaphase whereas HDAC3 remains on the chromatin during all mitosis stages. The sirtuins show a partial chromatin association at prophase and they are not seen at the chromatin from metaphase on. Finally, we found that some of the HATs (EP300 and HAT1) remain on the chromatin while others (CBP and GCN5L2) fall off. These results are only partially consistent with previous observations. The displacement of HDAC1 and HDAC2 from metaphase through the rest of mitosis stages was reported previously in MCF7 cells (He et al 2013). On the other hand, HDAC3 was found on mitotic chromatin in HeLa-S3 cells (Li et al 2006) but not on MEFs (Bhaskara et al 2008). Similarly, SIRT1 was found by us to evict the mitotic chromosomes at the metaphase stage in HeLa-S3 cells (**Fig. 3**), while in MEFs it seems to be retained on the chromatin along all mitotic stages (Fatoba & Okorokov 2011). These discrepancies are either due to the use of different techniques or due to the use of different experimental systems. Further research is required to determine whether the conflicting results reflect the above differences or variations between primary and transformed cells. Our conclusions are based on the use of small-molecule HDACs inhibitors that target specific HDACs or groups of histone deacetylases (**Table 1**). To mitigate the dependency on each of the small molecules’ specificity, we aimed to employ multiple inhibitors to target key deacetylases. Thus, we evaluated the involvement of HDAC2 by using MS-275, Chidamide and CAY10683 and studied HDAC3 by using RGFP966, Chidamide and MS-275 (**Table 1**). However, for SIRT1 we could only use EX-527, and thus, we currently cannot rule out the involvement of additional sirtuins in the process and further research is required to study this possibility.

Based on immunofluorescence and western blot analyses (**Figure 2**) we deduced that HDAC2 and SIRT1 are released from the mitotic chromosomes during metaphase. Previous studies (Festuccia et al 2019, Pallier et al 2003, Teves et al 2016) suggest that PFA fixation artificially causes such eviction. To evaluate this possibility, we repeated the IF experiments of SIRT1 and HDAC2 adding DSG crosslonking, since it was suggested that such crosslinking is better for studying mitotic retention of proteins (Festuccia et al 2019). In both cases we found similar results with the original PFA fixation and the DSG+PFA new fixation (**supplementary figure S7**). These results suggest that SIRT1 and HDAC2 are resistance to the artifacts of PFA on mitotic chromatin retention. This is consistent with the mild changes in H3K9ac levels upon inhibiting HDAC2 at metaphase (**Figure 2**).

Taken together, our results suggest a complex mechanism for H3K9 deacetylation during mitosis. It is carried out by a combination of HDACs that are activated at different mitotic stages and potentially via a reduction in HAT activity that is modulated by SIRT1. Additionally, we identified a temporal separation in the activity of the HDACs, with HDAC2 acting at prophase on most transcription-associated H3K9ac peaks, whereas HDAC3 contributes at a later stage. Further research is needed for studying the consequence of each wave of deacetylation on mitotic traits such as transcription repression and chromosome condensation. https://www.sciencedirect.com/science/article/pii/S0009898110004250?via=ihub

## Materials and methods

### Cell culture and cell cycle synchronization

HeLa-S3 cells and MEFs were grown in Dulbecco’s Modified Eagle’s Medium (DMEM) supplemented with 10% fetal bovine serum (FBS), penicillin-streptomycin, L-glutamine, sodium pyruvate, and 0.1% Pluronic F-68. Cells were incubated at 37°C and 5% CO_2_. DT40 cells were cultured in RPMI 1640 medium supplemented with 10% fetal calf serum, 1% chicken serum, 10 mM HEPES and 1% penicillin-streptomycin mixture at 39.5 °C with 5% CO_2_.

For STC mitotic arrest, cells were pre-synchronized in G1/S by addition of 2 mM thymidine for 16 h, washed with PBS, released for 3 h in fresh medium, and arrested with 10μM S-trityl-L-cysteine (STC) (164739, Sigma) for 12 h. For double thymidine synchronization, HeLa-S3 cells were grown on coverslips in Dulbecco’s Modified Eagle’s Medium (DMEM) supplemented with 10% foetal bovine serum (FBS), penicillin-streptomycin, L-glutamine, sodium pyruvate, and 0.1% Pluronic F-68. Cells were incubated at 37°C and 5% CO_2_. Cells were grown for 20 h, treated with 2 mM thymidine for 18 h, washed with PBS, released for 8 h in fresh medium, treated with 2 mM thymidine for 17 h, washed with PBS, released for 8.5 h in fresh medium, and harvested.

### HDAC and sirtuin inhibitors

The concentrations and specificities of the inhibitors we used are summarized in **Table 1**. The inhibitors were added for 8.5-9 hours (**Fig. 1a** and **c**).

### Chromatin isolation

Chromatin was isolated from STC-synchronized HeLa-S3 cell line using a previously published protocol (Sone et al 2002). Synchronized cells were collected by centrifugation and resuspended into a hypotonic solution of 75mM KCL (pH 5.7). After treatment of KCL hypotonic solution for 30 min, the cells were collected by centrifugation and resuspended into CAS buffer (0.1 M citric acid, 0.1 M sucrose, 0.5% Tween 20, pH 2.6). After lysis of the cell membrane in CAS buffer, the chromatin suspension was centrifuged at 190g for 3 min at 4°C. The chromatin-rich fraction was carefully recovered as supernatant and the nuclei rich fraction was recovered as the precipitation. The supernatant fraction was then centrifuged again at 1750g for 10 min at 4°C. The precipitated chromatin was resuspended in CAS buffer containing 0.1 mM phenylmethylsulfonyl fluoride. The isolated mitotic chromatin quantity was estimated by BCA method and stored in -80°C until further use.

Chromatin from asynchronous cells was isolated as described (Torrente et al 2011). Cells were resuspended in Buffer A (10 mM HEPES pH= 7.9, 10 mM KCl, 1.5 mM MgCl2, 0.34 M sucrose, 10% glycerol, inhibitor cocktail: 1 mM DTT, 0.5 mM 4-(2-aminoethyl) benzenesulfonyl fluoride hydrochloride and protease inhibitor cocktail). Triton X-100 was added to a final concentration of 0.1% and the suspension was incubated for 8 minutes on ice. The nuclear pellet was obtained by centrifugation (1,300g for 5 minutes at 4°C), washed with Buffer A and then resuspended in Buffer B (3 mM EDTA, 0.2 mM EGTA, and protease inhibitor cocktail) for 30 minutes on ice. The insoluble chromatin pellet was isolated by centrifugation (1,700g for 5 minutes at 4°C) and then resuspended in 15 mM Tris, pH= 7.5, 0.5% SDS. The isolated mitotic chromatin quantity was estimated by BCA method and stored in -80°C until further use.

### Immunoblotting

Isolated chromosomes were directly suspended and dissolved in SDS sample buffer (62.5 mM Tris-HCl (pH 6.8), 5% 2-mercaptoethanol, 20% glycerol, 2% SDS, and 0.005% bromophenol blue). Purified chromatin was separated by SDS-PAGE and transferred to 0.45 μm polyvinylidene difluoride membranes (Immobilon, Millipore, Merck KGaA, Darmstadt, Germany). Blots were incubated with primary antibodies HDAC1 (Cell Signalling # D5C6U; 1;1000), HDAC2 (Cell Signalling #D6S5P; 1:1000), HDAC3 (Cell Signalling #7G6C5; 1;1000), CBP(Cell Signalling #D6C5;1;1000),GCN5L2(Cell Signalling #C26A10; 1:1000),P300(Santa cruz #sc-48343; 1:100) HAT-1(Santa cruz # sc-390562; 1:1000), SIRT1(Cell Signalling #8469; 1;1000), SIRT2 (Santa cruz #sc-28298; 1:1000), SIRT6 (Abcam #ab62739; 1:1000) and SIRT7 (Santa cruz #135055; 1:1000), histone H3 acetyl K9 (Ac-H3K9; (Cell Signalling #C5B11; 1:1000). HRP-conjugated secondary antibodies (Jackson Immuno Research #111-035-003 and #115-035-003; 1:5000) were used. Immunoblots were developed with an ECL-plus kit. Equal loading of protein in each lane was verified by histone H3 (Cell Signalling #D1H2; 1:1000).

### Immunofluorescence staining

For immunofluorescence studies, HeLa-S3 cells and MEFs were grown on coverslips, fixed with 4% formaldehyde, (Thermo, Cat# 28908). Alternatively, HeLa-S3 cells were cross-linked with 2mM DSG (Sigma, Cat# 80424) for 50 minutes followed by 10 minutes incubation with 1% formaldehyde at room temperature (Festuccia et al 2019). After fixation cells were blocked in PBS containing 5% BSA and 0.1% Triton X-100. Cells were then incubated with a primary antibody - HDAC1 (Cell Signalling # D5C6U; 1:100), HDAC2 (Cell Signalling# D6S5P; 1:1000), HDAC3 (Cell Signalling #7G6C5; 1:100), CBP (Cell Signalling #D6C5; 1:100), GCN5L2 (Cell Signalling #C26A10; 1:100), P300 (Santa cruz #sc-48343; 1:100), HAT-1 (Santa cruz # sc-390562; 1:100), SIRT1 (Cell Signalling #8469; 1;100), SIRT2 (Santa cruz #sc-28298; 1:100), SIRT6 (Abcam # ab62739; 1:100) and SIRT7 (Santa cruz #135055; 1:100), H3K9ac (Cell Signalling #C5B11; 1:400) overnight at 4°C. After washing, cells were incubated with an anti-rabbit IgG or anti-mouse IgG conjugated to FITC probes (Invitrogen #A11034 and #A11001) for 1 h at room temperature (RT). After washing, slides were mounted with SlowFade Gold antifade reagent with DAPI (Sigma) and immunofluorescent signals were viewed using an Olympus fluorescence microscope using CellSens software. All images were taken under fixed scaling and were normalized to only secondary background control to avoid false significance due to overexposure. Images were analysed with ImageJ (Fiji version) to quantify mean intensity/pixel in the DAPI stained regions.

### *In Vitro* HAT Activity Assay

*In vitro* histone acetylation assays were performed on 25 μg of mitotic chromatin (isolated as described above and quantified using the bicinchoninic acid colorimetric assay system (Sigma)) in 50 mM Tris-HCL, 10% glycerol, 1mM DTT, 1mM PMSF, 50nM TSA, 0.1 mM EDTA supplemented with 100μM Acetyl-CoA and 10 μg His-tagged Histone H3 (BPS Bioscience #52023). The reaction was incubated at 37°c for 1 h. HAT activity was quantified by: i) a colorimetric assay quantifying the release of Co-A using DTNB (2-nitrobenzoic acid) (Sigma) by measuring the absorbance at 412nm; ii) isolating the His-tagged H3 by Ni-NTA column and measuring H3K9ac by immunoblots using anti-H3K9Ac antibody (Cell Signalling # C5B11; 1:1000). All acetylation assays were performed in duplicate.

### *In vitro* HDAC3 Activity Assay

Mitotic HeLa-S3 cells were lysed in cell lysis buffer (0.5% NP-40, 20 mM Tris-HCl (pH 7.4), 500 mM NaCl, 0.5 mM EGTA, 10% Glycerol, 0.5% Triton-X 100, complete protease inhibitor cocktail and 1 μM of zinc citrate.). Mitotic lysate was precleared by the protein A magnetic bead (Cell signalling) for 20 min at room temperature. After protein quantification by the bicinchoninic acid colorimetric assay system (Sigma), 1mg protein samples were used for HDAC3 immunoprecipitation using four microliters of HDAC3 antibodies (Cell Signalling) and incubated at 4 °C overnight, followed by 2 h incubation with 20 μL protein A magnetic beads. The captured beads were rinsed with lysis buffer five times and i) boiled in 5× SDS loading buffer for 5 min for confirming HDAC3 immunoprecipitation by immunoblot (after brief centrifugation, supernatants were loaded onto 12% SDS-PAGE gels and detected using rabbit-anti-HDAC3 antibody (1:1000; cell signalling) and HRP-conjugated secondary antibodies). Or ii) immunocomplex was eluted from bead in 0.2 M glycine pH 2.6 by incubating the sample for 10 minutes and neutralized by adding equal volume of Tris-HCL pH 8.0, and stored at -20°c for further use.

To perform HDAC activity assay, we used 50 ng of immunoprecipitated HDAC3 and 4μg of His-tagged H3 (BPS Bioscience #52023; acetylated by pre-treatment with whole cell extract and isolated by Ni-NTA column), in HDAC Assay buffer (10 mM Tris-HCl (pH 8.0), 10 mM NaCl, 1 μM ZnSO4 10% glycerol, and complete protease inhibitor mixture), incubated at 37°C for 1hrs. After the incubation, SDS-PAGE loading buffer is added to the reaction and runs on 12% SDS-PAGE. HDAC3 activity in the presence or absence of inhibitors was assessed by immunoblotting using anti H3K9ac antibody (Cell Signalling # C5B11; 1:1000).

### ChIP-seq

ChIP experiments were carried out as described in (Mikkelsen et al 2007, Texari et al 2021). All ChIP experiments were carried out on 2×10^5^ cells. 1.5×10^5^ HeLa-S3 cells (unsynchronized or STC synchronized with or without the indicated treatments) and 0.5×10^5^ unsynchronized and untreated DT-40 cells were cross-linked with 4% formaldehyde and mixed. The cell mixture was lysed and sonicated by Covaris to obtain an average chromatin size of 200-700 bp. Immunoprecipitation, using anti-H3K9ac (C5B11, Cell Signalling Technology), was carried out with inverting at 4°C for 14– 16 h. Antibody-chromatin complexes were pulled-down using Protein A/G magnetic beads (Dy-10001D and Dy-10003D, Invitrogen), washed and then eluted. After cross-link reversal and Proteinase K treatment, immunoprecipitated DNA was extracted with 1.8x Agencourt AMPure XP beads (BCA63881, Beckman Coulter).

### Library preparation and sequencing

ChIP DNA (or unenriched whole cell extract) were prepared for Illumina sequencing as described (Yehuda et al 2018). Briefly, DNA was subjected to a 50 μl end repair reaction, cleaned by 1.8× AMPure XP beads, followed by a 50μl A-tail reaction. The products were cleaned and ligated to 0.75 μM Illumina compatible forked indexed adapters. Ligation products were size selected in order to remove free adaptors. Ligation products were amplified with 15 (Input DNA) or 18 PCR cycles (ChIP DNA). Amplified DNA was size selected for 300–700 bp fragments using AMPure XP beads. The final quality of the library was assessed by Qubit and TapeStation. Libraries were pooled and sequenced on NextSeq (Illumina), using a paired-end protocol. The sequencing depth of each library is provided at **Supplementary Table S1**.

### Bioinformatic analysis

ChIP-seq reads were aligned to the human (hg38) or chicken (galGal6) genomes using Bowtie2 (Langmead & Salzberg 2012). All regions listed in the ENCODE hg38 blacklist, were excluded from all analyses. Duplicate alignments were removed with Picard Tools MarkDuplicates (http://broadinstitute.github.io/picard/). Only reads with mapping quality > 30 were used in the analysis. Peak detection on merged unsynchronized replicates (with a whole cell extract dataset as a control) was done using MACS2 (Zhang et al 2008) with default parameters and q-value of 0.05.

Metagene analyses were done using the python package Metaseq (Dale et al 2014). The metagenes were aligned to human transcription start sites (n=60580, taken form ENSEMBL GTF files), enhancers and insulators (n=4274 and 2025, respectively, taken from (Javasky et al 2018)); and to chicken TSS (n=7234, taken form ENSEMBL GTF files), enhancers (n= 20633, retrieved from the enhancer atlas (Gao & Qian 2020), and insulators (n=13510 CTCF binding sites, retrieved from GSM1253767). IGV snapshots show TDF files created from the aligned sequence data (using the count command from IGV tools (Robinson et al 2011, Thorvaldsdottir et al 2013). HOMER annotatePeaks.pl was used to quantify sample coverage at each peak, shown in **Supplementary Figure S6**. Quantification of read coverage for all analyses in all datasets were normalized to the total number of reads multiplied by a normalization factor derived from the chicken promoter data.

### Data availability

The Chip-Seq datasets produced in this study are available in Gene Expression Omnibus GSE168180. (reviewer token – wtsfwacqndmzjct)

## Acknowledgments

We thank the members of the Core Research Facility in the Hebrew University School of Medicine; Dr. Idit Shiff and Dr. Abed Nasereddin for generating genomic data. This work was supported by research grants from the Israel Science Foundation (grant # 1283/21 to I.S.), ISF-NSFC (grant #2555/16 to I.S.), The Binational Science Foundation (joint program with NSF; grant # 2019688) and NSF/MCB-BSF (grant # 2003358 to A.Go).

## Authors contributions

SG, RM, LH, AGr and MF performed the experiments. RM, RR and LH performed the bioinformatics analyses with supervision by AGo, CB and IS. SG, IS, AGo and SE designed the experiments. IS and AGo wrote the manuscript with input from all the other authors.

## Conflict of interest

The authors declare that they have no conflict of interest.

## Supplementary figures and table

**Figure S1.**
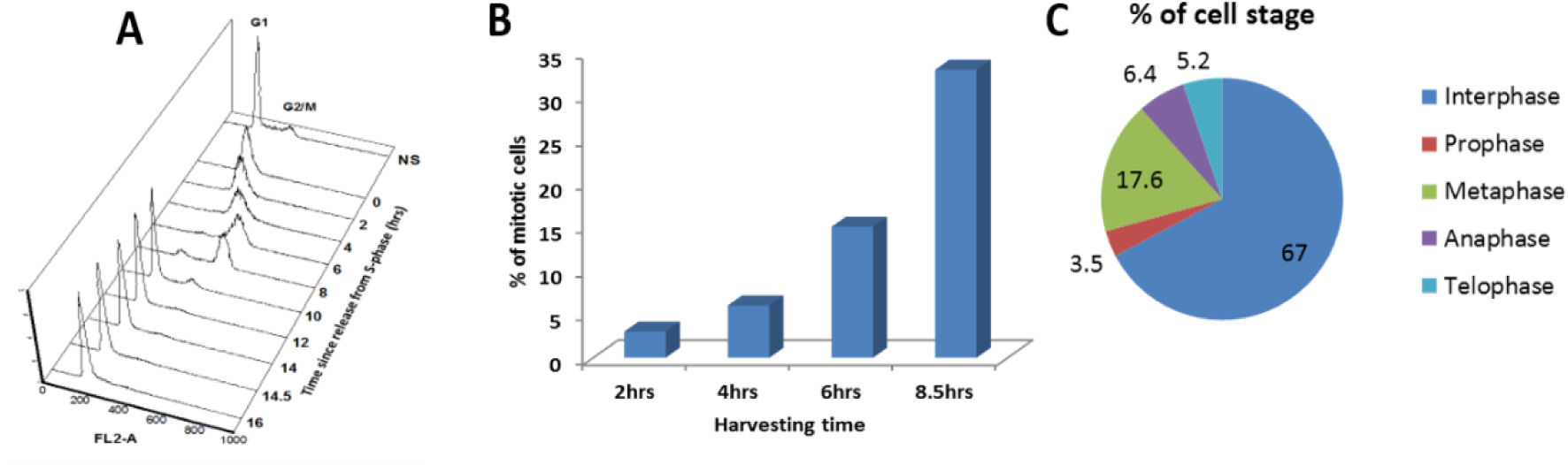
FACS analysis of double thymidine block (DTB) synchronized Hela S3: (A)The cell cycle distribution of cells synchronized by double thymidine block was monitored by FACS. The number of mitotic cells (arbitrary units) is plotted against DNA content for time points after release. (B) Bar graph showing the percentage of cells in mitosis at various time points after release from the DTB. (C) Pie chart showing the percentage of cells at different mitotic stages at the 8.5 hrs time point.

**Figure S2.**
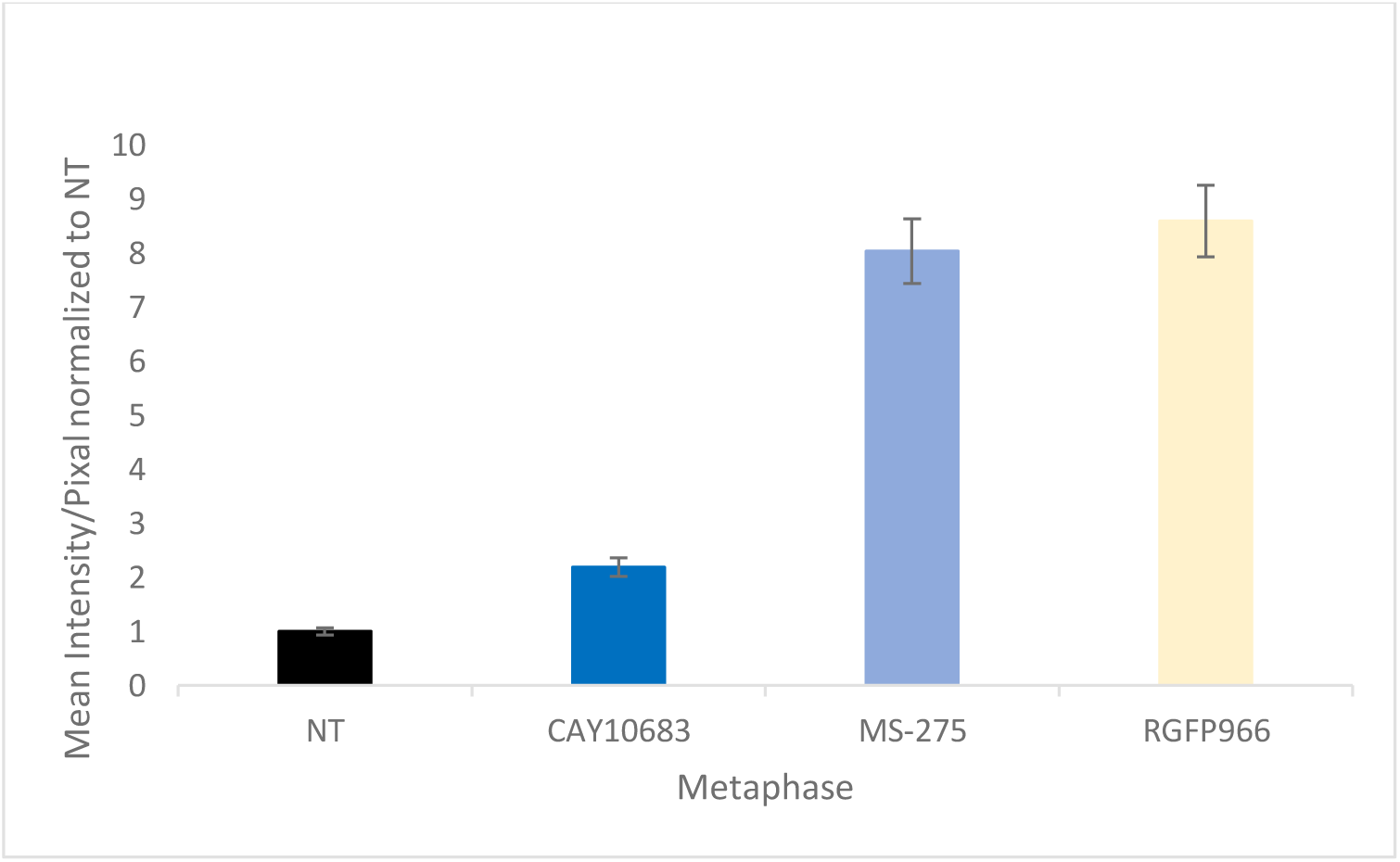
Quantification of the effects of key HDACi on H3K9ac levels on metaphase chromosomes. Quantification of the immunofluorescence results of HeLa-S3 cells enriched for mitotic stages stained for H3K9ac. Intensity values represent the mean ± SEM of 88, 128, 91 and 103 cells for NT, CAY, MS and RG treatments respectively. H3K9ac levels at metaphase chromsomes treated with either RG or MS are significantly different from both NT and CAY treated cells (P<10^−26^; two side ttest).

**Figure S3.**
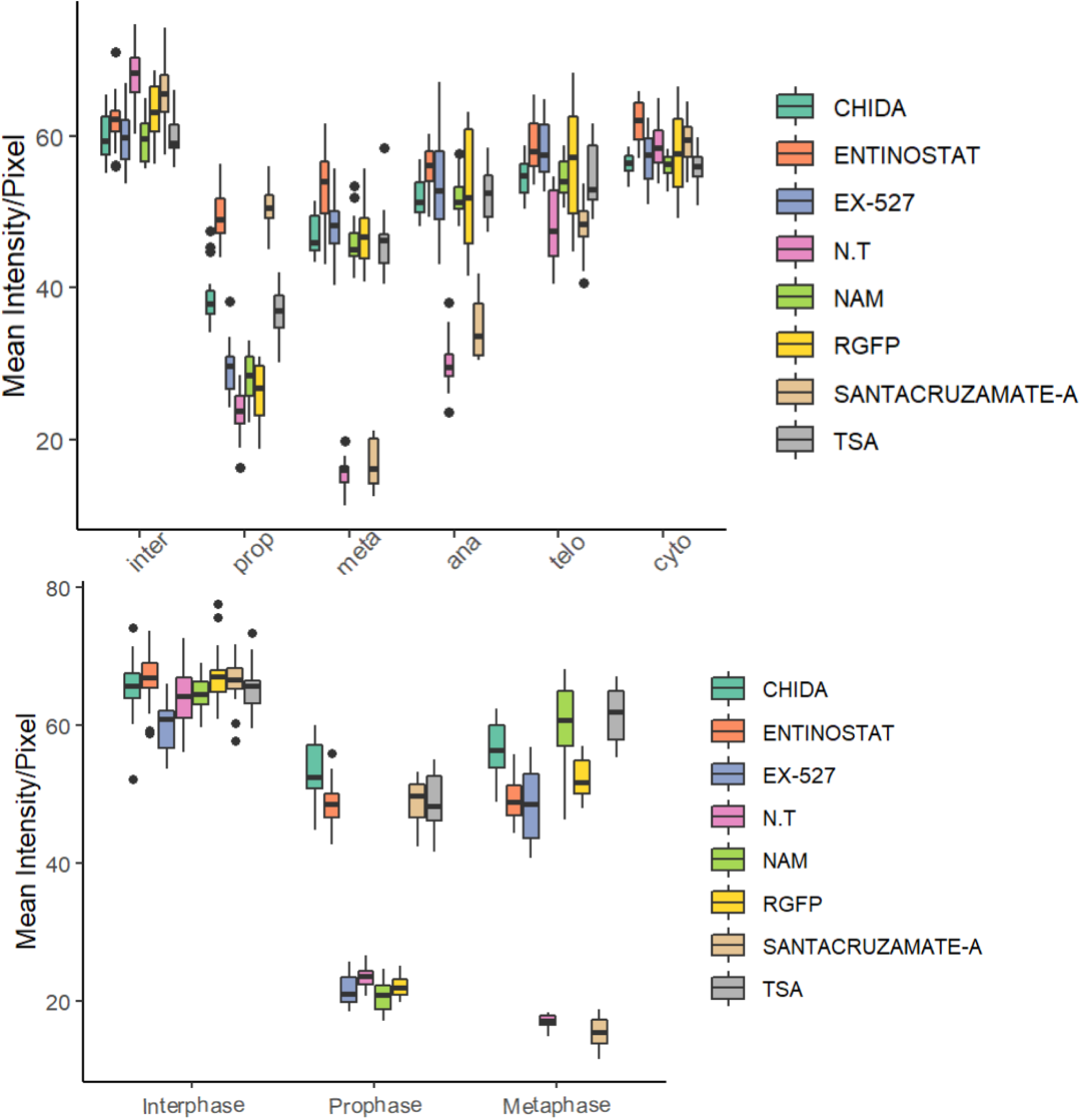
Box plot representation of the results of figure 2. For HeLa-S3 (up) and for MEF (bottom).

**Figure S4.**
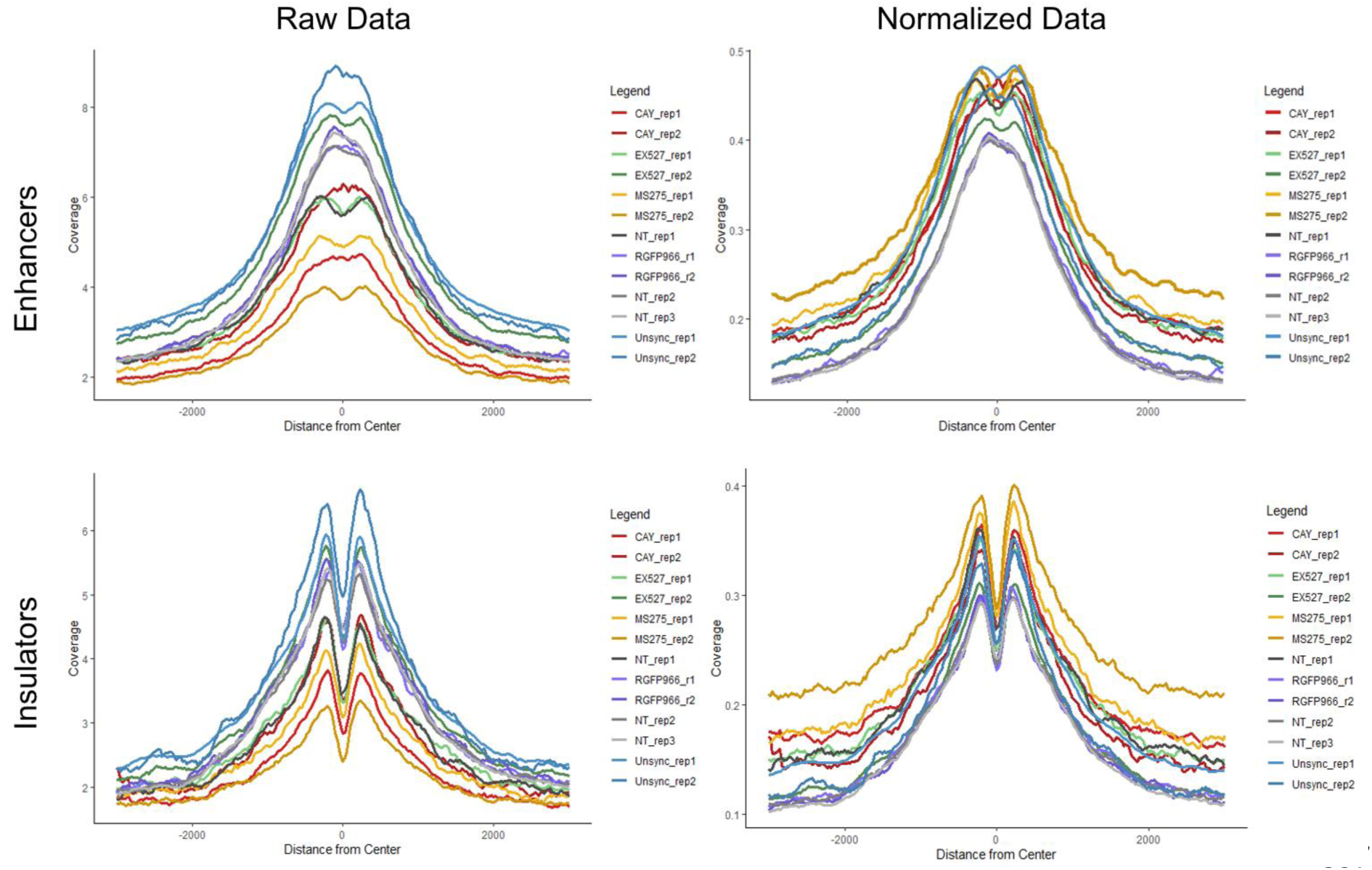
Metagene analyses of Chicken enhancers and insulators. Metagene plots showing H3K9ac enhancer and insulator occupancy in chicken DT-40 cells before (left) and after (right) normalization (according to promoter occupancy (see methods)(.

**Figure S5.**
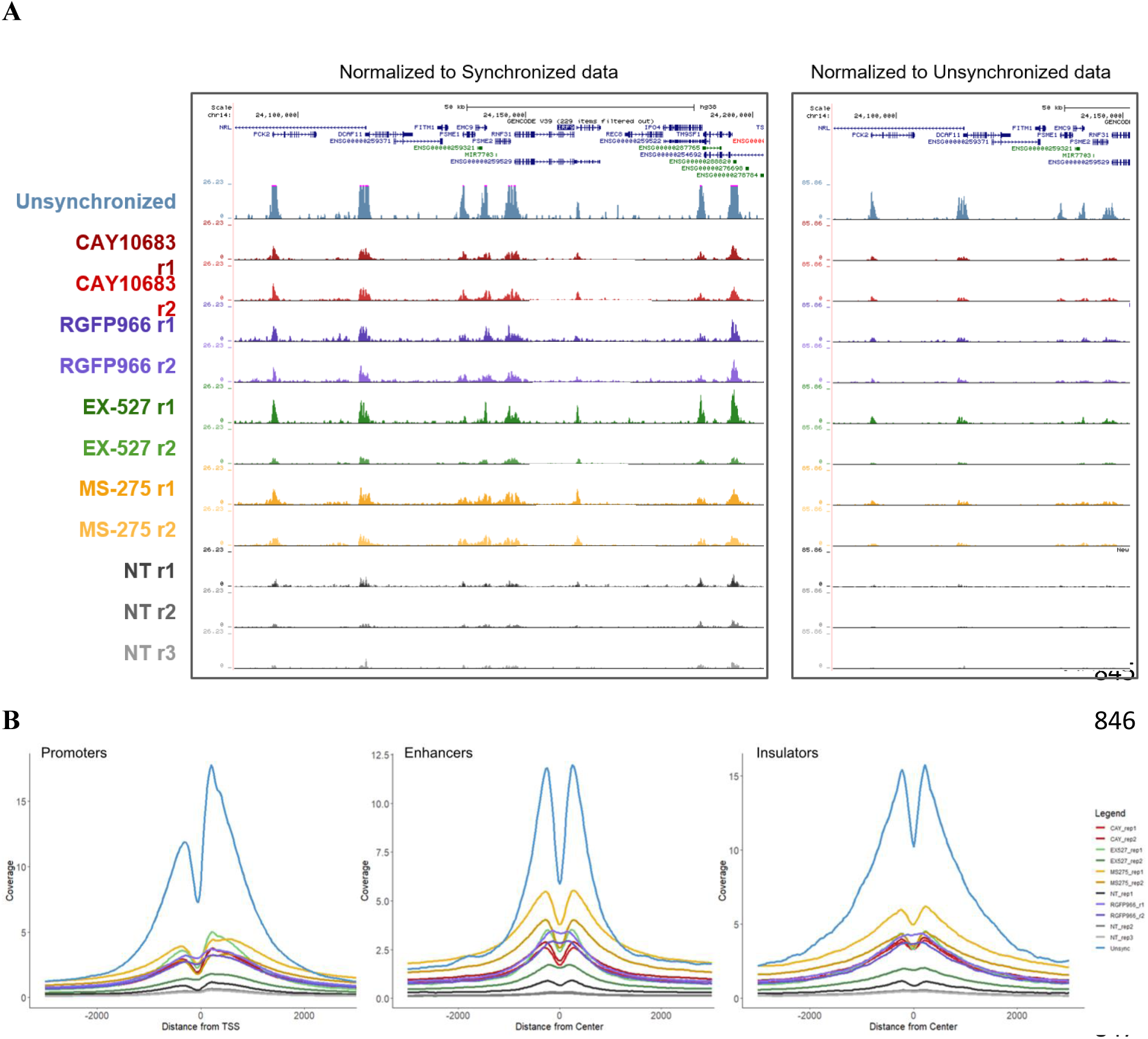
HDACs activity at transcriptionally associated regions (replicates) (A) Left shows UCSC representation of approximately 150kb region (Chr14:24,085,920-24,238,259) of H3K9ac ChIP-seq results under the indicated conditions. Data scaled to show inhibitor-treated peaks, Unsynchronized sample peaks out of range are highlighted in pink. Right shows a portion of the same genomic region but all samples are normalized equally. (B) Metagene plots showing H3K9ac occupancy around promoters, enhancers and insulators in HeLa-S3 cells. The data were normalized using the DT-40 promoter data.

**Figure S6.**
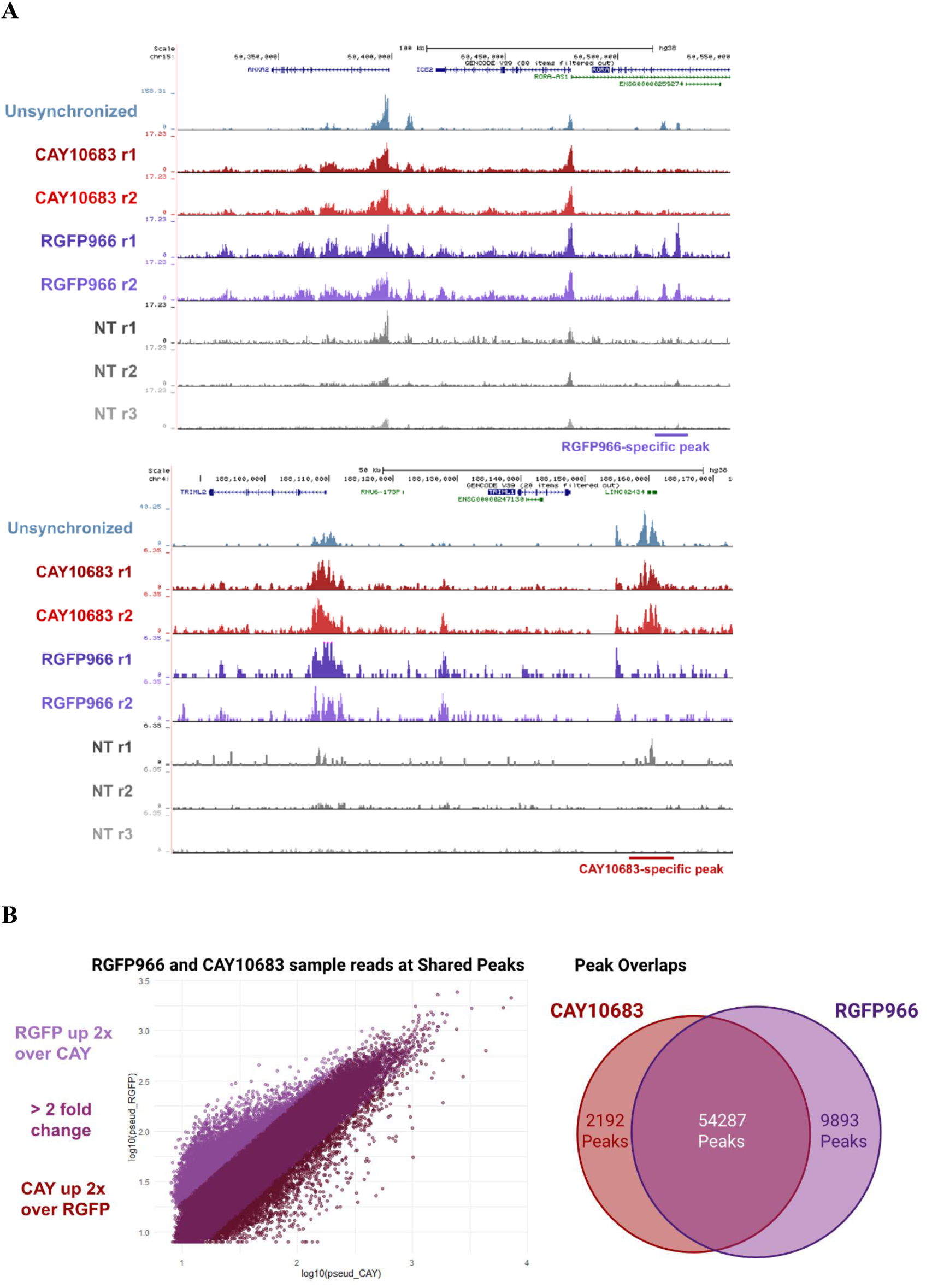
IGV plots of regions specifically effected by HDAC2 or HDAC3. (A) IGV representation of two genomic regions with RGFP966 and CAY10683 specific peaks. The locations of the differential peaks are indicated. (B) Scatter plot and Venn diagram depicting the differences between the H3K9c peaks in STC synchronized cells treated with either RGFP966 or CAY10683.

**Figure S7.**
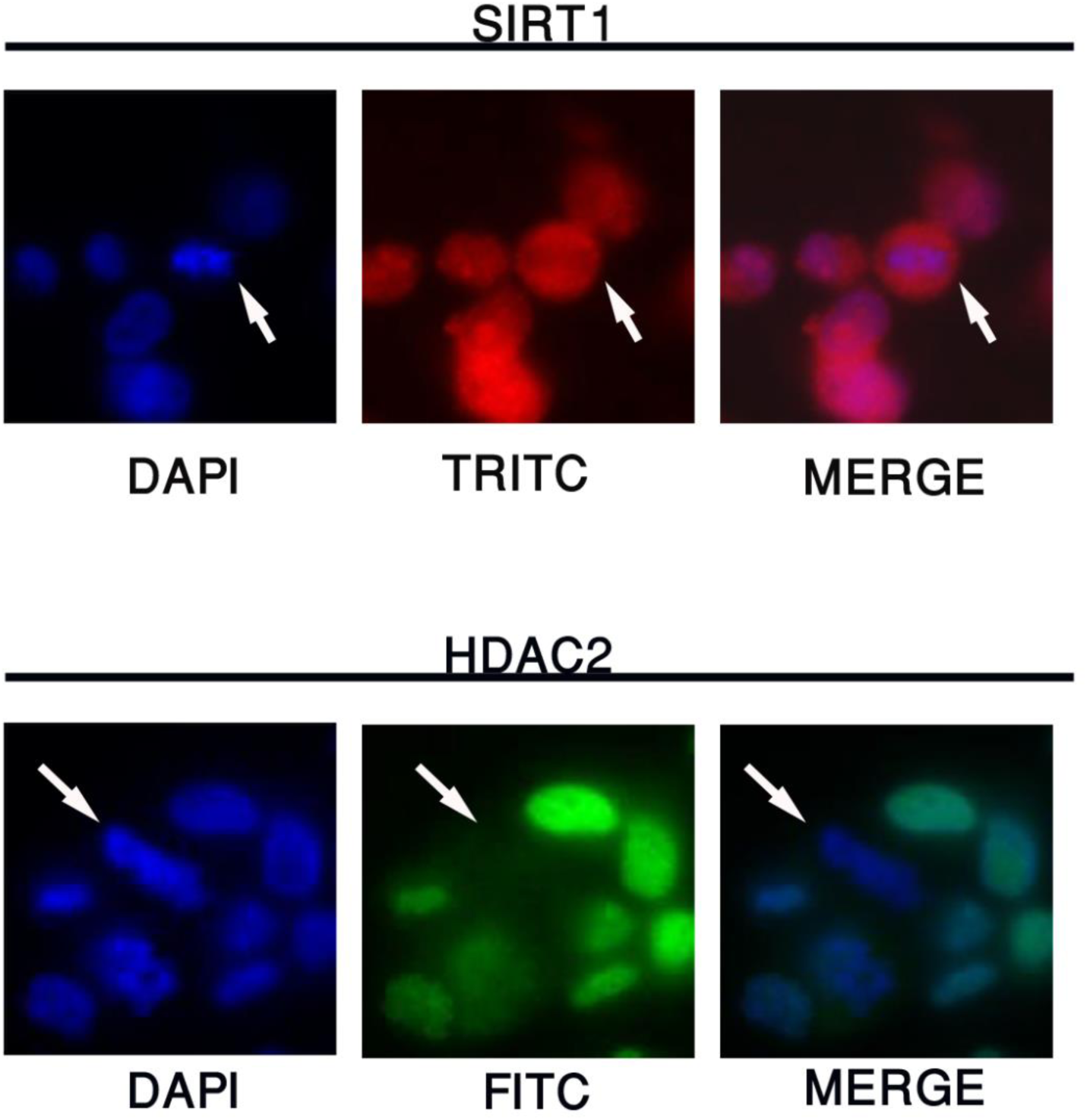
HDAC2 and SIRT1 are released from the mitotic chromatin. Immunofluorescence of HeLa-S3 cells cross-linked and fixed with DSG and FA. Arrows point to cells in metaphase. Note, that both SIRT1 and HDAC2 are depleted from the mitotic chromosomes.

**Table S1.** Sequencing depths and ChIP efficiency. <u>Normalization factor to correct for ChIP efficiency and read depth:</u> A factor used to account for differences in ChIP efficiency between samples. This factor was derived by the differences in the chicken promoter average occupancy and read depth.

| Sample name | number of reads | Normalization factor to correct for ChIP efficiency and read depth |
| --- | --- | --- |
| **Repeat-1 (Figure 5)** |  |  |
| Unsync_H3K9ac_rep1 | 25735442 | 0.629828 |
| Sync_NT_H3K9ac_rep1 | 5761776 | 1.4703905 |
| Sync_RGFP966_H3K9ac_rep1 | 10447729 | 2.747743 |
| Sync_CAY10683_H3K9ac_rep1 | 54414614 | 1.085428 |
| Sync_EX527_H3K9ac_rep1 | 10774458 | 1.222262 |
| Sync_MS275_H3K9ac_rep1 | 70298372 | 2.258310 |
| **Repeat-2 (Figure S5)** |  |  |
| Sync_NT_H3K9ac_rep2 | 9199420 | 0.554521 |
| Sync_NT_H3K9ac_rep3 | 10635598 | 0.485372 |
| Sync_RGFP966_H3K9ac_rep2 | 11245186 | 2.523053 |
| Sync_CAY10683_H3K9ac_rep2 | 68982790 | 1.480612 |
| Sync_EX527_H3K9ac_rep2 | 26117748 | 0.704151 |
| Sync_MS275_H3K9ac_rep2 | 69855678 | 1.652898 |

## Notes

### Competing Interest Statement

The authors have declared no competing interest.

### Summary of Updates

In response to reviewer requirements we 1. explored the effect of cross linking. 2. increase the number of measurements. 3. Added statistics

